# The circadian system is affected by Alzheimer’s disease independently from amyloid beta deposits

**DOI:** 10.64898/2026.08.25.744599

**Authors:** Hugo Calligaro, Brian Khov, Karina Noel, Aidan Glina, Laura van Rosmalen, Ramesh Ramasamy, Yushi Li, Michael Tun Yin Lam, Hiep Le, Keun-Young Kim, Won-Kyu Ju, Mark Ellisman, Satchidananda Panda

## Abstract

Circadian disruption, notably sleep disturbances, serves as an early indicator of Alzheimer’s disease (AD), preceding cognitive symptoms like memory loss. The suprachiasmatic nucleus (SCN) governs biological rhythms and receives direct retinal input via melanopsin-expressing retinal ganglion cells (mRGCs) to synchronize with environmental light cycles. The anatomical and functional basis for circadian disruption in AD remains unclear. Here, we explored the multi-level relationships between gene expression, the SCN connectome, and regulations of sleep and circadian rhythms in the APP/PS1 mouse model.

The sleep architecture of APP/PS1 mice displayed significantly reduced rapid eye movement sleep (REM), associated with a reduced daily core body temperature amplitude and locomotor hyperactivity. Lastly, APP/PS1 mice showed an impaired response to acute light pulse stimulation and present hyperactivity of mRGCs at a young age and hypoactivity of these cells at older ages. These physiological functions are known to be, at least in part, regulated by the SCN, the main target of mRGCs.

We noted several modifications in SCN connectomics using serial blockface electron microscopy (SBEM), including a reduction of the dendro-dendritic chemical synapse (DDCS) network that receives a large part of the retinal input and is thought to be crucial for synchronicity between SCN neurons. In addition, we observed multiple signs of dystrophy, including modifications of the shape of dendrites and cell soma, accumulation of aggregated lysosomes, and swelling of axons. At the same time, we investigated the changes in gene expression using spatial transcriptomics. The SCN presents changes in the expression of genes associated with synapse formation, cell adhesion, and neurite growth.

These results suggest that, despite the absence of amyloid plaques in the ventral hypothalamus, the SCN of APP/PS1 mice still undergo profound gene expression changes, impacting connectomics and physiological functions.

**Graphical abstract:** 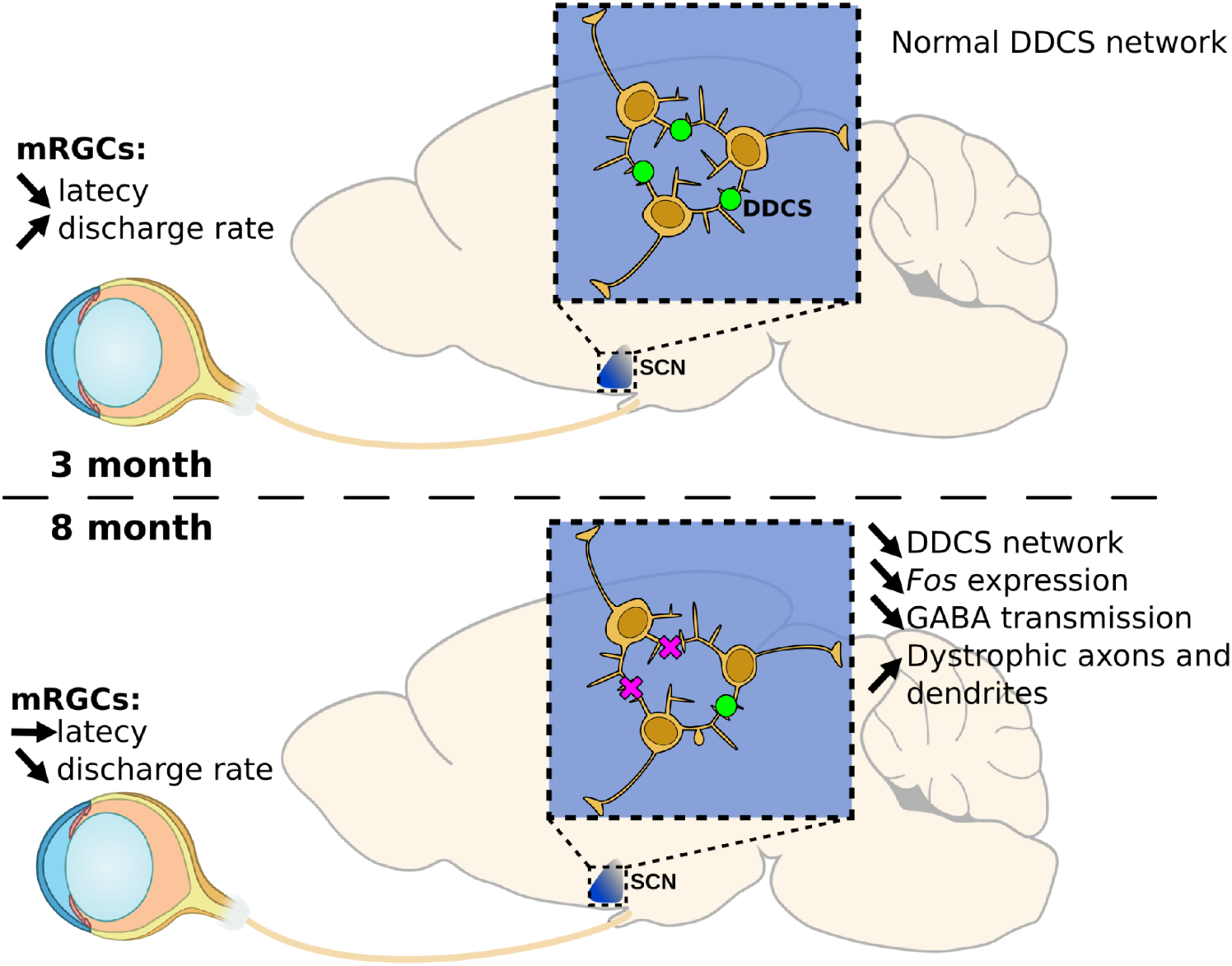

## Introduction

Alzheimer’s disease (AD), a subset of dementia, is an increasing concern in the world’s population as it represents ∼60-70% of all dementia cases and affected 7 million people in the United States in 2025, according to the World Health Organization. Individuals with this disease often present with cognitive impairments, including difficulties with performing daily activities and recalling memories. Current knowledge hypothesizes that accumulation of amyloid-beta and tau proteins drives the progression of AD and is thus of interest in treatments (Lane et al., 2018; Weller and Budson, 2018). Despite having potential therapeutic options, many clinical trials have not observed success, likely attributed to the advanced progression of AD. As a result, early detection methods are crucial for treating this disease (Phan and Malkani, 2019). Studies have observed individuals with AD exhibit symptoms of circadian dysfunction, such as the deterioration of rhythmicity in the sleep/wake cycle and psychiatric symptoms during sunset hours, which is known as “sundowning” (Leng et al., 2019; Phan and Malkani, 2019; Volicer et al., 2001). Sleep disruption is seen in the preclinical stage of AD, suggesting that sleep could be a potential biomarker. Notably, when amyloid-beta protein levels were measured, it was associated with poorer sleep quality, suggesting AD has a circadian basis (Ju et al., 2013; Mattis and Sehgal, 2016).

Changes in rhythmicity draw attention to the suprachiasmatic nucleus (SCN), the central regulator of all behavioural circadian rhythmicity. This brain region receives direct input from the light environment through a subset of retinal ganglion cells, the melanopsin-expressing retinal ganglion cells (mRGCs) (Evans, 2016; Hatori et al., 2008; Kim et al., 2019). The rhythmic expression of genes in SCN neurons results in coordination of the rhythms in our bodies, such as melatonin, temperature, and cortisol (Evans, 2016; Leng et al., 2019; Van Erum et al., 2018). Disruptions in all of these biomarkers are associated with AD. A microscopic look into the SCN reveals deterioration and neuronal loss, which may contribute to dysregulated circadian rhythms (Phan and Malkani, 2019; Van Erum et al., 2018). Additionally, studies have observed mRGC loss as well as alterations to dendritic morphology and size (La Morgia et al., 2023, 2017, 2016). Although the cellular basis of the SCN’s role in AD remains largely unknown, it is suspected that the failure to transmit information from the light environment appropriately may explain symptoms of circadian disruption (Esquiva et al., 2017; La Morgia et al., 2016; Phan and Malkani, 2019).

Here, we sought to identify the cellular and molecular basis of the observed circadian perturbations by studying circadian-related functions with a multi-scale approach in an AD mouse model. We thus evaluated the light response of mRGCs and established the SCN connectome at two crucial ages to estimate disease progression. Then, we measured sleep regulation and the modification of the transcriptome in the brain at a large scale using, respectively, state-of-the-art polygraphic recordings and image-based single-cell spatial transcriptomics. We found that, despite the absence of amyloid beta deposits, we observed significant changes in the gene expression and synaptic organization of SCN neurons at an early stage of the AD progression.

## Methods

### Animals

All animal care and procedures were approved by the Institutional Animal Care and Use Committee (IACUC) of the Salk Institute for Biological Studies and in accordance with ARRIVE guidelines. We used male APPswe/PS1ΔE9 mice and WT littermates in each experiment. All mice are maintained in a 12/12 light/dark cycle at 400 lux (White LED light) with food and water ad libitum. The mouse strain used for this research project, B6C3-Tg(APPswe, PSEN1dE9)85Dbo/Mmjax, RRID: MMRRC_034829-JAX, was obtained from the Mutant Mouse Resource and Research Center (MMRRC) at The Jackson Laboratory, an NIH-funded strain repository, and was donated to the MMRRC by David Borchelt, Ph.D., McKnight Brain Institute, University of Florida.

### Functional assays

#### Sleep recording

To measure sleep regulation, a telemetric transmitter (DSI, HD-X02) was implanted in the abdominal cavity of WT and APP/PS1 mice to record continuous electroencephalogram (EEG) and electromyogram (EMG) data in addition to core body temperature and locomotor activity. Specifically, two EEG biopotential electrodes were implanted in contact with the dura at the top (1mm anterior and 1mm left lateral to Bregma) and back of the skull (3 mm posterior and 3mm right lateral to Bregma, contralateral to the first lead). For EMG, the activity of the dorsal neck muscle was measured by placing two biopotential leads within the same bundle of the trapezius muscle. These data were captured through a telemetry receiver (DSI, RPC-1) placed underneath the mouse cage. The receiver is connected to the computer through a Data Exchange Matrix with a sampling rate of 300Hz using the Dataquest ART acquisition software (DSI, Minnesota, USA).

Traditional sleep assessment methods in mice, relying on the manual and visual interpretation of EEG (electroencephalogram) and EMG (electromyogram) waveforms, inherently carry risks of subjectivity and bias. To address these limitations, we have developed an automated sleep scoring system grounded in deep learning technology. Our system is anchored by a two-layer neural network designed to accurately distinguish between wakefulness, rapid eye movement (REM) sleep, and non-REM (NREM) sleep stages in mice, based on EEG and EMG data. The network was trained using manually annotated data representing wake and sleep states, with input features derived from the normalized root-mean-square of extracted EMG signals, periodogram of total EEG power (0.5-25Hz), and the power ratio of the five frequency bands – Delta (0.5-3.9Hz), Theta (4-7.9Hz), Alpha (8.0-12.0Hz), Sigma (12-16Hz), and Beta (16-24Hz) – over the total EEG power using Discrete Fourier Transform. This method effectively leverages the nuanced variations in frequency and amplitude that typify different sleep and wake states. To refine and validate our model, we employed a dataset comprising 94,839 10-second epochs for training, supplemented by 23,661 epochs for validation (20%), achieving an accuracy rate of 91.45%.

NREM sleep epochs were categorized using k-means clustering based on six normalized EEG features: total power, Delta, Theta, Alpha, Sigma, and Beta power ratios. We analyzed 166,661 10-second epochs from six male mice, setting k = 6 to generate six clusters (Lam et al, in preparation). Cluster validation was performed by examining distinct EEG feature distributions and their temporal patterns over the 24-hour period. The clusters were labeled NREM clusters 1–6, based on their temporal phase from ZT 0. This dataset served as the discovery cohort, and new recordings were then clustered by assigning each epoch to the nearest centroid from the initial clusters, using Euclidean distance.

#### Light phase-shift experiment

Singly housed male WT and APP (n = 5 for each genotype) were first entrained in a 12L/12D cycle for 20 days in a running-wheel cage. Subsequently, animals were maintained in constant darkness (DD) to examine the free-running period, calculated by periodogram analysis using ClockLab software (Actimetrics). Phase shifts were studied using a single 30-min monochromatic light pulse (470 nm) at 1e14 photons/cm²/s applied at circadian time 16 (CT16), 4 hours after lights off. At the time of the stimulation, mice were individually transferred to a cage equipped with a light stimulation frame surrounding the cage to obtain a homogeneous stimulation. After the light pulse, animals were returned to their home cages, and activity was monitored in DD for an additional 15 days. The magnitude of a light-induced phase shift was determined from the difference between the regression lines of the activity onsets before and after the light stimulation, extrapolated to the day following the light pulse. The transient responses on the 3-4 days immediately after the pulse were discounted (Pittendrigh and Daan, 1976).

#### Multielectrode Array recording

Retina recordings were done as described previously (Calligaro et al., 2026; Mure et al., 2019). After a dark-adapted mouse was euthanized, we isolated the retina under dim red light and maintained it in oxygenated artificial cerebrospinal fluid (aCSF) in the dark. Patches of retinas of APP/PS1 mice were mounted on a 256-electrode multielectrode array (MEA, Multichannel Systems, Reutlingen, Germany) with ganglion cells facing down and continuously perfused with oxygenated aCSF at 34°C. To isolate RGCs from rods’ and cones’ response to light, the retinas were perfused with aCSF supplemented with a synaptic blocker cocktail containing D-AP5 (50µM), CNQX (20µM), and L-AP4 (50µM) for at least an hour before any exposure to light. Negative thresholds for spike detection were set at 5 times the standard deviation (SD) of the noise on each channel. Spike cut-outs, consisting of 1 ms preceding and 2 ms after a suprathreshold event, along with a timestamp of the trigger, were written to the hard disk. Recorded retinas were exposed to full-field light stimulation of increasing irradiance (5e11 photons/cm²/s to 1e15 photons/cm²/s) using monochromatic LEDs (470 nm, LuxeonStar 5, luxeonstar.com). Irradiances and durations of the light stimulations were controlled by custom Python scripts.

For each channel, spike cut-outs were sorted into trains of a single cell using Offline Sorter (Plexon, Denton, TX). Data analysis and visualization were performed using Neuroexplorer (Plexon) and custom R scripts (Rproject). Different parameters of the response of the cell to light were determined. The response discharge rate was calculated as the average of the action potential discharge during the response duration, minus the average discharge rate 30s before the start of the light stimulation (baseline). The duration of the response was defined as the time during which the discharge rate was over a threshold defined as the baseline + 2SD. The response was considered over when the discharge rate was below the threshold for 3 consecutive seconds. To consider that a cell responds to light stimulation, the duration of its response should last longer than 1s.

### Connectomics

#### Tissue preparation

Two APP/PS1 mice of 3 and 8 months (respectively 3M and 8M) were used in the connectomics experiment. Mice were anesthetized with an intraperitoneal injection of ketamine/xylazine and transcardially perfused with Ringer’s solution containing heparin and xylocaine (∼1 min flush), followed by approximately 50 ml of 0.1% glutaraldehyde/4% prilled paraformaldehyde in 1× PBS. Brains were dissected and post-fixed in 4% formaldehyde in PBS on ice for 2 h, then sectioned into 100-µm-thick slices. The core regions of the SCN and OPN were identified by histological markers under a dissection microscope and prepared for serial blockface scanning electron microscopy (SBEM) as previously described (Calligaro et al., 2023; Kim et al., 2019). Sections containing the regions of interest were washed in 0.15 M sodium cacodylate buffer and processed for SBEM using an osmium-based staining protocol. Briefly, sections were incubated in 2% OsO₄/1.5% potassium ferrocyanide/2 mM CaCl₂ in 0.15 M sodium cacodylate for 1 h, washed in water, then transferred to filtered 0.05% thiocarbohydrazide for 30 min. Sections were washed again in water and incubated in 2% aqueous OsO₄ for 30 min. After a final water rinse, sections were transferred to filtered 2% aqueous uranyl acetate overnight at 4°C. Sections were then washed with ddH₂O at room temperature and stained en bloc with 0.05% lead aspartate for 30 min at 60°C. Following staining, sections were washed with water and dehydrated on ice through a graded ethanol series (50, 70, 90, 100, and 100%) for 10 min at each step, then washed twice with dry acetone. Sections were infiltrated in 1:1 Durcupan ACM:acetone overnight, transferred to 100% Durcupan resin overnight, and flat-embedded between mold-release-coated glass slides. Resin was cured at 60°C for 72 h. SBEM volumes were acquired at a 2.5-kV accelerating voltage with a raster size of 15,000 × 15,000 or 20,000 × 20,000 pixels, a pixel dwell time of 1 µs, a pixel size of 4 or 5 nm, and a section thickness of 50 nm. Prior to each acquisition, a low-magnification (500×) image of the block face was collected to confirm anatomical location based on tissue landmarks. Following acquisition, image intensity drift across the volume stack was corrected by histogram normalization in Digital Micrograph; files were then converted to MRC format, scaled to 8-bit, and manually traced for volumetric reconstruction and analysis. One SBEM volume of the SCN and one of the OPN were obtained from each mouse.

#### Reconstruction and analysis

To analyze these datasets, we used the publicly available software package IMOD, specifically developed for the visualization and analysis of EM datasets in three dimensions (Kremer et al., 1996); http://bio3d.colorado.edu/imod/). Cross-sectional contours were manually traced for consecutive data slices in the z dimension to determine the boundaries of user-defined objects. For some objects, contours were traced in every other data slice in the z dimension. These contour profiles were used for three-dimensional volumetric reconstruction of the cell body, axons, boutons, and organelles.

### Manual segmentation

#### Counting of cells

To count the neurons, astrocytes, and vascular cells in each dataset, a five-by-five grid was created. The entirety of the dataset was scanned systematically by starting from one corner and going through all of the created subvolumes and marking each element of interest with a different color. This process was done until all the volume was covered.

#### Random selection of axons and dendrites

To estimate the changes in the connectome, we randomly selected 100 dendrites and 100 axons using the criteria listed below on an 11 x 11 grid from the middle of the volume. The dendrite and the axon closest to each intersection were marked, and then fully skeletonized every five z-steps. If two of the randomly selected neurites ended up connecting after skeletonization, we selected an additional element to obtain a final population of 100 of each neurite.

#### Identification of axons, dendrites, and boutons

To discriminate between axons and dendrites, we looked for characteristic morphological features (See Supplementary Video 1 for examples). Dendrites usually have a constant, large diameter with dendritic spines and dendritic intrusions and mostly form postsynaptic elements of synapses. The axons have a smaller diameter with varicosities forming synapses, receive dendritic intrusions, and are systematically the presynaptic element of synapses. For the identification of boutons, we looked for areas of the axon that appear to be swollen to a diameter at least twice as large as the average diameter of the axon. Our criteria for swelling to be considered a synaptic bouton included the presence of at least one synapse identified as synaptic vesicles close to the cell membrane in contact with a postsynaptic element. This postsynaptic element can be a direct contact with a neighboring cell soma or a dendritic intrusion.

#### Volume quantification

To calculate the volume of organelles, nuclei, and spines, we fully segmented them individually every two z-steps. Their volumes were then extracted using *imodinfo*.

#### Synaptic input quantification

Synaptic sites were identified by scanning the neurites previously skeletonized and finding areas with synaptic vesicles close to the cell membrane and at least one of the following criteria: proximity between the presynaptic and postsynaptic element cell membranes and presence of dendritic intrusions. Synapses are then classified between axodendritic chemical synapse (ADCS) and dendrodendritic chemical synapse (DDCS) by identifying both elements using the specific criteria listed before. To avoid overestimation and underestimation of the density of synapses on dendrites, we excluded from the analysis all dendrites shorter than 2 mm.

#### Training and quality control

All persons involved in manual tracing were first trained to identify and label specific structures (axons, dendrites, soma) and ultrastructures (mitochondria, synapses, stigmoid bodies, etc.). All data were systematically checked at least once by a different person from the one who initially produced it.

### Spatial transcriptomics

#### Tissue collection and processing

Three 10-month old males of each genotype (WT and APP) were used in this experiment. Animals were euthanized using cervical dislocation at zeitgeber time 5 (ZT5), 5 hours after light on. Whole brains were extracted from the skull within 5 minutes and frozen in isopentane on dry ice for 1 minute. Subsequently, brains were embedded in Tissue-Tek OCT compound (Sakura Finetek, Torrance, CA, USA) by transferring the sample into isopentane surrounded by dry ice for 1 minute. Frozen embedded brains were then stored in air-tight bags at −80°C until cryosectioning. Coronal brain sections (10 µm) were prepared using a Leica CM1950 cryostat set at −18°C. Two sets of samples have been prepared: 6 coronal sections centered around the SCN (Bregma −0.34 mm; R0) and 6 coronal sections every 700 µm to obtain a large panel of brain regions (respectively −1.58, −2.30, and −3.08 mm; R1). During sectioning and collection on Xenium slides, one sample from R0 was damaged. We processed it through the Xenium protocol nonetheless. Post-run examination of its data parameters (cell number, transcript quality, and transcript number) was similar to other sections of the run; we thus decided to include it in the analysis.

All 12 coronal sections were placed onto 4 Xenium slides (PN 3000941, 10x Genomics, Pleasanton, CA, USA) according to the Xenium tissue preparation guide (10x Genomics, CG000579). The Xenium mouse brain gene expression panel (10x Genomics, PN 2000825) was used to detect gene expression levels of 247 genes for cell type identification. We prepared the Xenium slides following 10X Xenium guidelines (10x Genomics, #CG000581). Briefly, frozen sections were then fixed in 4% formaldehyde to retain RNA and subsequently permeabilized in methanol to allow RNA to be accessible. Probes were hybridized at 50°C overnight, followed by ligation and primer hybridization for rolling circle amplification according to the Xenium gene expression protocol (10x Genomics, CG000582). Subsequently, autofluorescence quenching and nuclei staining were performed. Sections R1 were additionally processed with the multi-modal cell segmentation staining to label (10X Genomics, CG000749) the multi-modal cell segmentation targets ATP1A1, E-Cadherin, and CD45 (segmenting cell membrane boundaries), 18S ribosomal RNA (cytoplasm), and α-smooth muscle actin (SMA)/Vimentin (interior protein staining). Tissue sections were then imaged using the automated Xenium analyzer (10x Genomics, CG000584). Fluorescent probes were hybridized and imaged following probe removal for a total of 15 cycles. The intensity is measured in 750 nm Z-stack slices in 4 different color channels to measure target gene expression levels.

The resulting data were analyzed using Xenium Onboard Analysis version 1.7.1.0 (R0) and 2.0.0.10 (R1). Despite being processed with the same reagents and containing a similar number of cells, R1 sections resulted in a larger number of transcripts compared to R0 sections. We thus processed each experiment separately as described below.

#### Amyloid beta immunostaining

Immunostaining was done for the R1 sections after completing the Xenium Analyzer run. Sections following the section of R0 samples were collected during tissue sectioning and immunostained for amyloid-β. Slides were first permeabilized and blocked in 0.3% Triton X-100 and 1% BSA in PBS for 1 hour. Subsequently, slides were incubated in primary anti-β-Amyloid antibody (6E10 monoclonal, 1:500, BioLegend, 803004) overnight at 4°C. After washing in PBS, sections were incubated in secondary antibodies (Alexa Fluor 488, 1:200, Invitrogen, A-11001) at room temperature for 2 hours. Both antibodies were diluted with the blocking solution. Nuclei were counterstained with DRAQ5 (1:1000, ThermoFisherScientific, 62254). Then, coverslips were mounted with an antifade mounting medium (Vectashield, H-1700). Sections were imaged with Axioscan 7 Microscope Slide Scanner at 20x magnification.

#### H&E staining

Following the Xenium run (R0) or the immunostaining imaging (R1), the sections were processed for hematoxylin and eosin staining following Post-Xenium Analyzer H&E Staining protocol (CG000613). Cover slips of R1 samples were removed by incubation for 30 minutes in PBS at room temperature. Briefly, the sections were incubated in fresh 10 mM Sodium Hydrosulfite for 10 minutes. The sections were then incubated in hematoxylin (Sigma Aldrich, MHS16) for 20 min, after which the sections were washed and dehydrated through increasing concentrations of ethanol (70% and 95%). The sections were then immersed in eosin (Leica, 3801615) for 2 min. The sections were then dehydrated with 95% and 100% ethanol and washed in Xylene for 6 minutes. Finally, coverslips were mounted with Permount Mounting Medium (Electron Microscopy Science, 17986-05). Sections were imaged with an Axioscan 7 Microscope Slide Scanner.

#### Cell segmentation

Cell segmentation for sections R1 was done by the Xenium onboard software using the multimodal cell segmentation labels (Supplementary Figure 1A). Cells in sections R0 were labeled with 4′,6-diamidino-2-phenylindole (DAPI), and the Xenium Analyzer onboard cell segmentation using DAPI only consists of the detection of the nuclei and a soma expansion (15 µm). The cell segmentation of R0 by soma expansion results in a large overestimation of the soma size and the inclusion of neuropils within their soma, which could lead to transcripts being incorrectly assigned to cells. Using the DAPI staining obtained during the Xenium processing, we resegmented R0 cells using Qupath (v.0.5.1) cell detection and soma expansion (5 µm), and re-assigned transcripts, which resulted in cell parameters (cell area, number of unique genes, total transcripts) comparable to those of cells in R1.

### Data Analysis

#### Data preprocessing

Data was preprocessed using Scanpy (Wolf et al., 2018). We imported single-cell gene expression data from 10x h5 files. Samples from the same experiment were processed together. Transcripts with q-values ≥ 20 were included in the analysis. We filtered out all cells with fewer than 40 total transcripts or fewer than 5 unique genes. The gene expression is then normalized and log-transformed by cells using scanpy.pp.normalize_total and scanpy.pp.log1p methods before further analysis.

#### Clustering and annotation

To define the cell types present in our dataset, we employed the Leiden algorithm (resolution = 2) to define clusters. To annotate the resulting clusters, we correlated them with single-cell annotations obtained with MapMyCells (Allen Institute), which correlates single cells with a reference taxonomy of the whole mouse brain (CCN20230722) (Yao et al., 2023). Clusters presenting multiple cell classes were subclustered using the Leiden algorithm (resolution = 0.2). Similarly, to investigate subtypes of microglia and SCN neurons, we subsetted the population of interest and re-runned the clustering analysis using the Leiden algorithm (resolution = 0.2). Resulting clusters were considered as subtypes when presenting different gene markers and/or spatial organization.

#### Brain region

The brain is highly organized into brain regions containing different neuronal populations, sometimes without clear anatomical landmarks. To obtain an objective and reproducible delimitation of brain regions, we developed a method using the cell type annotation to define the brain regions. First, cell types belonging to the same regions were grouped together (e.g. excitatory and inhibitory neurons of the cortex were merged into a “cortical neuron” cluster). Cluster information was then used to establish contiguous regions of distinct cell types with a multi-step computational pipeline. First, annotated cells were grouped using a K-Nearest Neighbors (KNN) classifier (k = 20) to mitigate noise arising from mixed cell populations in spatial proximity. For each cell type, a graph was constructed where nodes represented individual cells, connected by edges to their nearest neighbors. To ensure the preservation of local cell neighborhoods, the graph was refined by applying a Minimum Spanning Tree (MST) and pruning edges with lengths exceeding two standard deviations from the mean. Small disconnected subgraphs with fewer than 100 cells were further removed to isolate robust contiguous regions. The spatial data were then subdivided into a grid (50 x 50 µm), and for each grid cell, the predominant cell type was assigned based on the majority class within the corresponding chunk. Grid cells without a clear majority were labeled as background. To ensure smooth and contiguous boundaries, empty grid cells were filled by propagating neighboring cell type assignments. This produced non-overlapping, contiguous regions corresponding to distinct cell type areas. Finally, these regions were converted into polygon representations, enabling further downstream spatial and biological analyses.

#### Gene expression analysis

We used the Wilcoxon signed-rank test (scanpy.tl.rank_genes_groups), ties-corrected, to identify differentially expressed genes (DEG). Obtained p-values are corrected by the Benjamini-Hochberg method. We established the DEG between WT and APP samples for each condition studied (cell types, brain regions, unassigned transcripts). Only genes expressed in at least 10% of the cells of each condition with a mean normalized expression over 0.2 were considered in the analysis. Normalized gene expression was averaged for each condition, and the fold change was calculated as FC = APP/WT, then transformed by a base 2 logarithm. All genes with adjusted p-value under 0.05 and FC over 1.2 (= log2FC over 0.26) were considered significantly different.

#### Plaque detection

To detect and segment automatically amyloid plaques, we trained a pixel classifier using a Random Tree algorithm using a set of 20 manually annotated plaques as training data (Qupath, v. 0.5.1). Objects were then created with a minimal area of 400 µm² and exported as polygons to be aligned with the Xenium dataset. The majority of plaques were observed in the cortex, corpus callosum, amygdala, and hippocampus in brain sections of the APP/PS1 mice. No plaques were detected in WT mice brain sections. To evaluate the gene markers associated with amyloid plaques, we compared the gene expression of all cells within the plaques with the cells of the same brain region outside of the cells using the method described previously.

#### Unassigned transcripts

To evaluate the role of transcripts outside of cell somas, we extracted all gene transcripts with the “Unassigned” cell label and with q-values ≥ 20 from the raw Xenium files and mapped them to a 50 x 50 µm grid. The grid is then mapped to the sample borders on the same coordinate system to accurately calculate the area of each grid cell within sample borders. Grid cells with an area under one-tenth of the maximal area were removed to avoid under- and over-estimation of the transcript density. The resulting matrix is transformed into an annotated dataframe and analyzed similarly to the cells.

### Statistical analysis

Results are expressed as means +/- Standard Error of the Mean (SEM). All statistical analyses were performed using R (R project). Comparisons of the two groups were done using a nonparametric bootstrap rank-sum test (Mann–Whitney U test). Comparisons of multiple groups were done by using ANOVA on ranks (Kruskal–Wallis H test) followed by a bootstrap post hoc test (pairwise Mann–Whitney U tests). A statistically significant difference was assumed with a p-value inferior to 0.05.

### Data and code availability

The SBEM image volumes are available on Cell Image Library (XX). The image-based spatial transcriptomic dataset generated for this study is available on the GEO database (GSE328654). All custom scripts generated in this study are available on GitHub (https://github.com/Carboneinerte).

## Results

### 1. Sleep regulation of REM and NREM sleep is altered in APP/PS1 mice

A set of WT and APP/PS1 mice (n=4 for each genotype) were implanted with DSI probes to record EEG, EMG, locomotor activity, and core body temperature across time to evaluate sleep regulation in Alzheimer’s Disease. We recorded these parameters in two 4-day sessions for a total of eight days of recording per mouse. The mice were maintained in a cycle of 12 hours of light (400 lux of white L.E.D.) and 12 hours of darkness. The continuous recording was extracted in 10-second bins and automatically annotated using a specialized algorithm (see Methods for details). The resulting data were then averaged for each Zeitgeber Time (ZT), with ZT0 corresponding to light on.

Both the WT and APP/PS1 mice exhibit daily variations in wakefulness, NREM sleep, REM sleep (Figure 1A), and core body temperature. We observed that the APP/PS1 mice have a REM sleep reduction (p < 0.01, Figure 1A). However, no significant difference in total sleep is observed (p = 0.14). To analyze the architecture of NREM sleep in more detail, we employed a recently developed clustering method (Lam et al.) based on unsupervised clustering to identify six subtypes composed of varying ratios of brainwaves. These clusters have reproducible daily patterns (Figure 1C). Subtype 1, in particular, is linearly decreasing during the resting phase, then increases during the active phase. The APP/PS1 mice have a similar pattern but with differences in their distributions. Indeed, we observed that, in APP/PS1 mice, subtype 1, which has the highest ratio of low frequency bandwidth (Delta), is reduced during the light phase (p<0.01, Figure 1B,C). In contrast, subtype 2, which is composed of high frequency bandwidth (Alpha-Sigma), is inversely increased (p<0.01, Figure 1B,C). Subtype 2 is also increased during the active phase in APP/PS1 mice.

**Figure 1:**
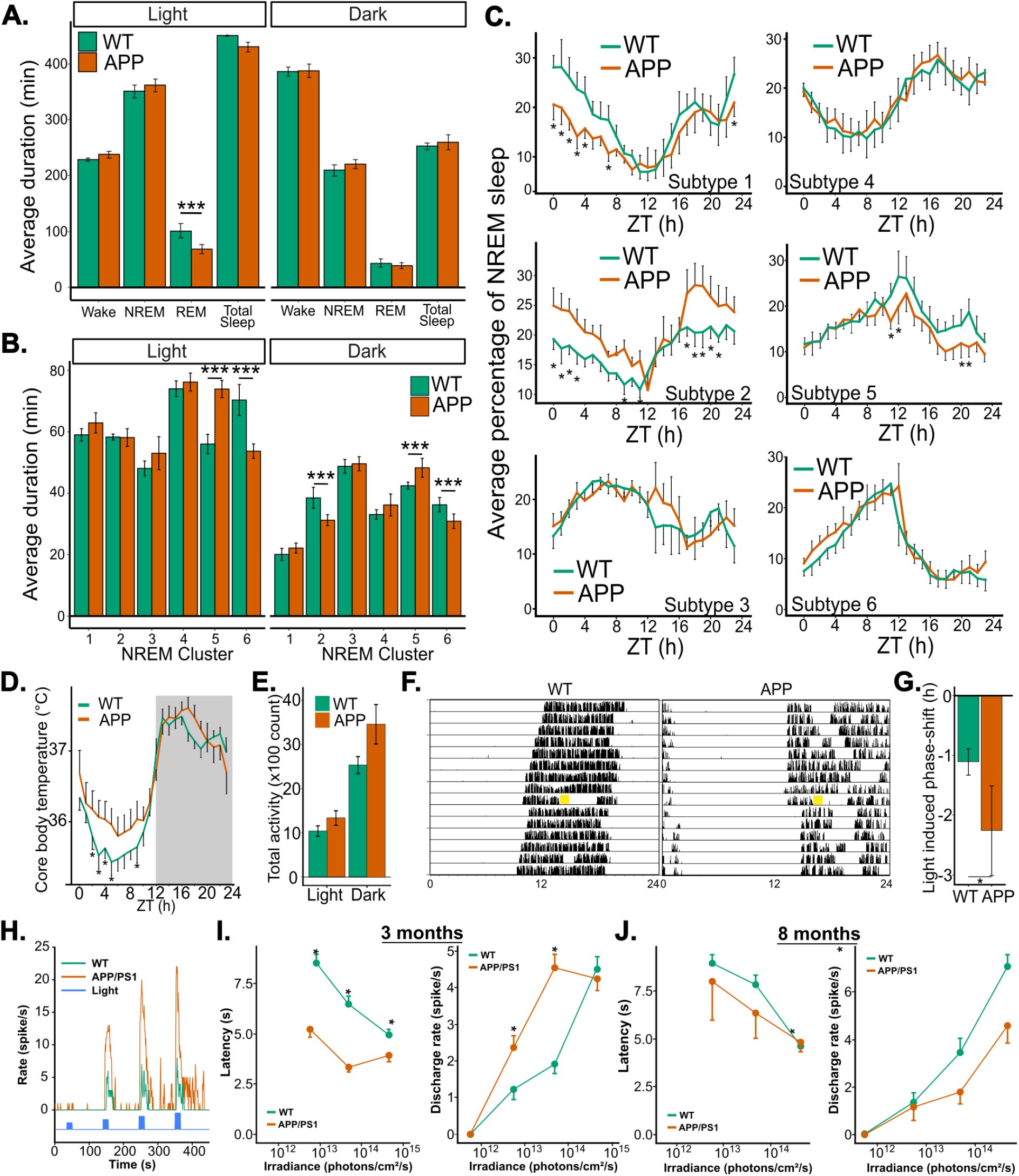
APP/PS1 mice present reduced REM sleep and a change in NREM sleep architecture A. Average duration of wakefulness, NREM sleep, REM sleep, and total sleep in APP/PS1 mice and WT littermates during light and dark phases. B. Average duration of the NREM sleep subtypes in APP/PS1 mice and WT littermates during light and dark phases. C. Daily distribution of all six subtypes of NREM sleep. D. Daily changes of core body temperature averaged every 1h. E. Average locomotor activity of WT and APP mice in light and dark phases. F. Representative actogrammes of the wheel activity in constant darkness of WT and APP mice. The yellow squares represent the 30 min light pulse (470 nm, 1e14 photons/cm²/s) done at CT16. G. Phase shift induced at CT16 by 30 min of blue light stimulation (470 nm, 1e14 photons/cm²/s). H. Visualization of the light protocol used to assess the electrical response to light of melanopsin cells (blue line) and representative examples of the light response at 3 months. I-J. Quantification of the latency (left) and the discharge rate (right) of the electrical response to light of melanopsin cells at three (I) and eight (J) months. Results are expressed as mean +/- SEM. n = 4 for each genotype. Statistical analysis was done using One-Way ANOVA. * = p-value < 0.05.

These results demonstrate an effect of AD on sleep regulation and architecture. Multiple brain regions regulate these processes, but also depend on the SCN, which has yet to be examined in the context of AD. In addition, APP mice showed a reduced amplitude of their core body temperature due to a higher core body temperature during the active phase (Figure 1D) and a slight increase of their locomotor activity during the dark phase (WT = −1.11 ± 0.22; APP = −2.26 ± 0.75; p < 0.05; Figure 1E). The changes in sleep and temperature regulation were mainly observed during the light phase, or near the transition between dark and light, which raised the possibility of a change in the ability to receive and process photic information required to maintain the entrainment of rhythmic functions to the external environment. A common method for evaluating circadian functions is through the measurement of rhythmic locomotor activity and its response to light challenges. We maintained singly-housed WT and APP mice in special cages, recording their wheel activity. The mice were then placed in constant darkness for 10 days and subsequently exposed to 30 minutes of bright blue light (1e14 photons/cm²/s) at circadian time 16 (CT16), 4 hours after light off, a time known to induce maximal phase delay. APP/PS1 mice showed similar endogenous periods to their WT littermates (WT = 23.59 ± 0.08 h; APP = 23.85 ± 0.08 h; p = 0.07), but the light pulse induced a larger phase delay (WT = 1.11 ± 0.22 h; APP = 2.26 ± 0.76 h; p < 0.05; Figure 1G). Then, we tested the electrical response of melanopsin cells (mRGCs) to monochromatic light stimulation of increasing intensity (Figure 1H). Interestingly, at the pre-symptomatic age of 3 months, the mRGCs of APP mice presented a short latency associated with a discharge rate reaching its maximum with lower light intensity, compared to WT mRGCs (Figure 1I). This hyperactivity is, however, lost by 8 months. Indeed, at that age, we did not observe any latency differences between WT and APP, and a lower discharge rate at the maximal intensity (Figure 1J). Taken together, these results are converging to suggest that AD affects the circadian system, leading to impaired circadian regulation of sleep and activity, as well as an impaired response to acute and chronic modifications of the environmental light cycle.

### 2. AD affects all cell types across the brain

To examine the changes in gene expression induced by AD on the SCN, but also in the rest of the brain at a large scale, we used an imaging-based spatial transcriptomic technique (Xenium, 10x Genomics) with a 247-gene panel specifically designed for mouse brain samples. We sampled four different brain sections of each genotype and obtained a total of 1.7 million cells and 600 million transcripts. All 247 genes presented a quantity of transcripts superior to the noise level estimated by the negative control probes present in the panel. We then clustered and annotated the dataset using the MapMyCell database (Allen Brain Institute) based on Yao and colleagues’ (Yao et al., 2023) published dataset (Figure S1). We then confirmed the individual cell types by extracting, for each cluster, the top differentially expressed genes and comparing them to known markers of cell types. Using the Allen Brain Institute gene expression atlas as a reference, we identified 84 unique cell types (Figure 2A-B) across 23 brain regions.

**Figure 2.**
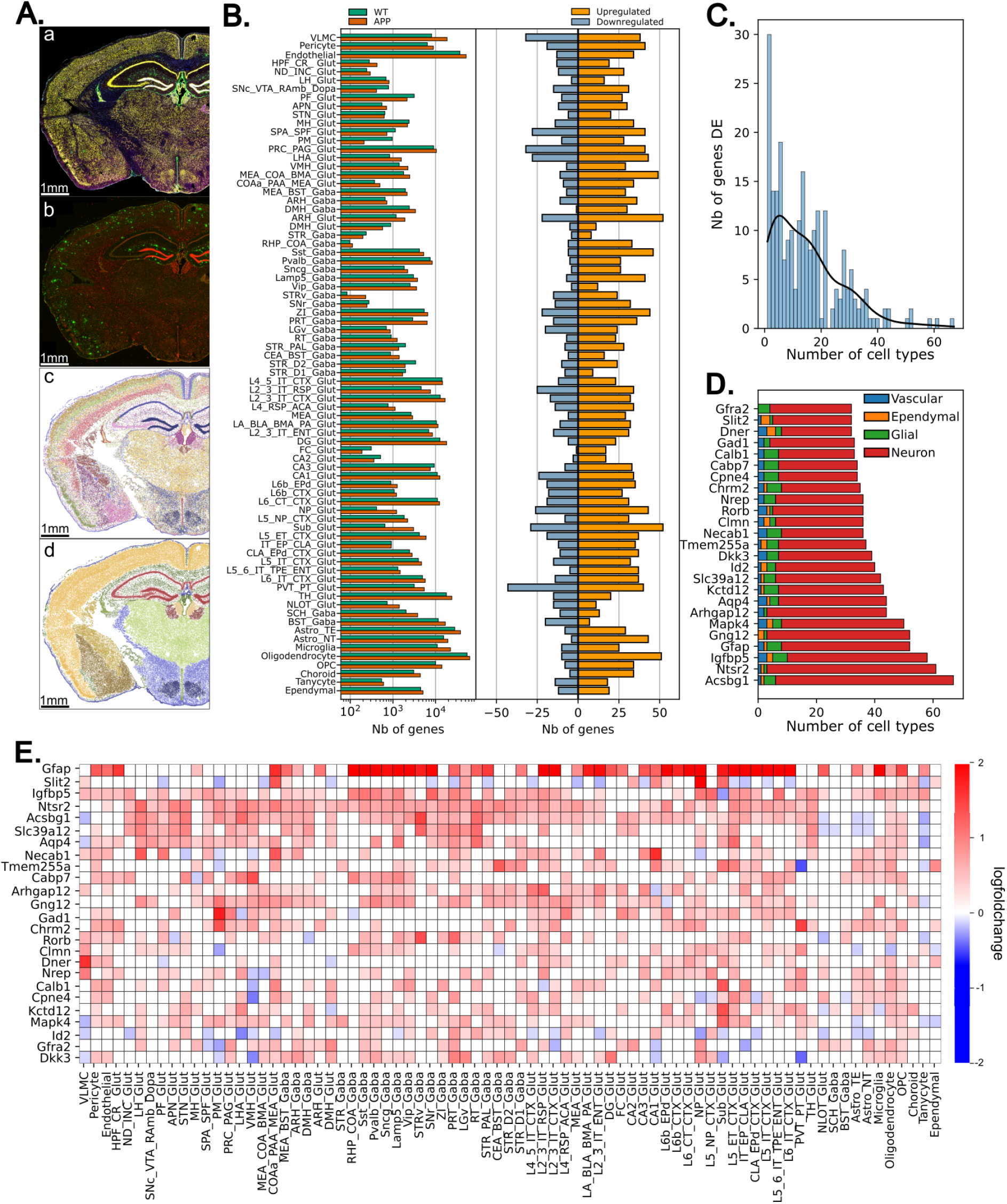
Cell type populations and differentially expressed genes. A. Representative examples of a brain section processed with spatial transcriptomics showing the cell segmentation markers (a; blue = DAPI, green = SMA/Vimentin, yellow=18S), the amyloid plaque immunostaining (b; red = DRAQ5, green = b-Amyloid), the cell types identified (c), and the brain regions (d). B. Cell number (left) and differentially expressed genes (right) for each cell type identified. C. Distribution of the number of cell types in which each gene is differentially expressed. D. Top 20 of genes differentially expressed in the most number of cell types. E. Heatmap of the logfold change of these genes in all cell types.

All cell types with balanced cell population between WT and APP genotypes were used for differential expressed genes (DEG) analysis. All of them presented at least one DEG, with a slight bias toward upregulation in APP mice (Figure 2B). If many of the DEG are specific to one or small number of cell types, we observed that some genes are changing in the majority of the cell types (Figure 2C-D). These DEG are distributed across all cell classes identified (neuronal, glial, ependymal and vascular cells) and are mainly upregulated (Figure 2E). *Gfap*, a marker for astrocytes and whose overexpression is one of the first signs of neuroinflammation, is highly upregulated, in particular in cortical and hippocampal regions. Other widely upregulated genes are associated with various functions, including neurotransmitter and synapse regulation (*Gad1*, *Chrm2*, *Slc39a12*, *Arhgap12*), immune functions (*Acsbg1*, *Ntsr2*, *Gng12*). The only gene mainly downregulated in APP, *Slit2*, is known to regulate axon guidance and migration. Interestingly, only three of these genes (*Gfra2*, *Kctd12*, *Slc39a12*) are significantly changed in SCN neurons.

### 3. Unassigned transcripts show an increase of inflammation markers in the cortex and hippocampus, but not in the hypothalamus

One specificity of image-based spatial transcriptomics is that it gives access to mRNA localized outside of the cell soma. This is crucial information in brain tissues, where an important part of translation occurs in neurites and glial processes, often in the proximity of synapses to ensure fast and reliable recycling of proteins. This can be informative of the changes happening outside of the cell soma (Figure 3A). We thus isolated the transcripts localized outside of cells (unassigned transcripts or UT), which range from 15% of all transcripts of a gene (*Col6a1*) to over 80% (*Unc3c*, Figure 3B). Interestingly, the genes which have a UT ratio over 50% are associated with modulation of trans-synaptic signaling (*Unc13c*, *Calb1*), neurotransmitter transport, and vesicle-mediated transport (*Angpt1*, *Syt2*).

**Figure 3:**
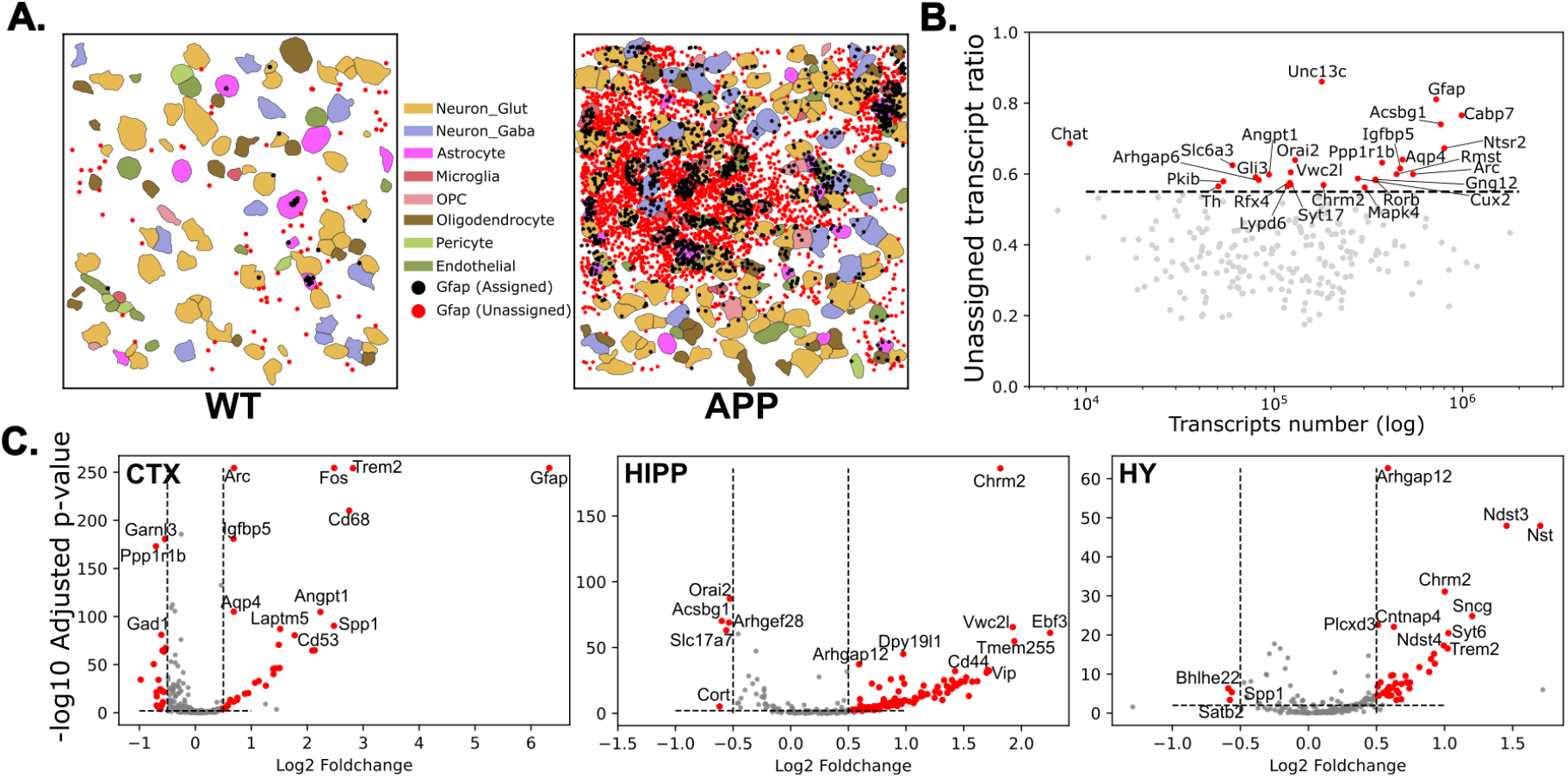
Unassigned transcripts show region specific changes between WT and APP samples. A. Examples of the distribution of Gfap assigned (black dots) and unassigned transcripts (red dots) in the cortex of WT (left) APP (right) samples. B. Ratio of unassigned transcripts over the total number of transcripts for each gene. Red dots correspond to genes with more than 50% of transcripts unassigned to a cell. C. Differently expressed genes in unassigned transcripts in APP cortex (left), hippocampus (middle), and hypothalamus (right). Red dots correspond to genes which are considered significantly different.

To allow us to compare the UT expression between conditions, we binned the UT into a 50 x 50 µm grid with each square of the grid treated as a pseudo-cell. The resulting dataset was processed similarly to the cells as described previously. We then calculated the differentially expressed UT in the cortex, hippocampus and hypothalamus. We observed an overexpression of inflammation-related genes (e.g. *Gfap*, *Trem2*, *Spp1*) specifically in the cortex (Figure 3C, left). In the hippocampus, the largest changes in gene expression were genes involved in cell proliferation (e.g., *Ebf3*, *Vwc2l*) and synaptic regulation (e.g, *Orai2*, *Chrm2*, *Vip*) (Figure 3C, middle). In the hypothalamus, UT related to synaptic organization (*Nst*, *Chrm2*, *Sncg*, *Cntnap4*) and metabolism (*Ndst3*, *Ndst4*, *Plcxd3*) are overexpressed in APP mice (Figure 3C, right). Non-synaptic cell communication is also affected with an increased expression of *Gjb2* (connexin 26), which supports glial cell communication, and a decrease of *Gjc3* (connexin 29), which is involved in myelin sheet maintenance by oligodendrocytes. Of the over 150 UT differentially expressed in these three regions, only 9 of them are common, suggesting that different mechanisms might be involved. In addition, some UT do not follow the same pattern in all regions. For example, *Spp1,* known to increase synapse pruning by microglia (De Schepper et al., 2023), is upregulated in the cortex and the hippocampus but downregulated in the hypothalamus. These results confirm that unassigned transcripts carry valuable information on the effect of AD on gene expression and the local variations between brain regions which suggest that different processes might be involved.

### 4. Identification of plaque-specific changes in gene expression

To establish the markers of AD within our panel of genes, we turned toward amyloid plaque, which are formed by amyloid-β oligomer aggregates. The staining for amyloid-β peptides (6E10) revealed amyloid plaques only in APP mice, distributed throughout the cortex, hippocampus, and amygdala. Rare signals were observed in the thalamic regions (Supplementary Figure 2). The plaques correlate with AlphaSMA/Vimentin staining used in Xenium cell segmentation and known markers of reactive astrocytes (Figure 4A). By comparing gene expression across all cell types outside and within plaques, we observed an upregulation of markers associated with the immune system (IL-1, IGF regulation, neutrophil degranulation), while downregulated genes corresponded to cellular development and differentiation, synaptic activity, and cell junctions organization (Figure 4B). When examining, in the cortex, the cell types present within the boundaries of the plaques, we observed a large increase in the density of microglia, but not of astrocytes, compared to the outside of plaques (Figure 4C). This suggests a specific recruitment of microglia to the amyloid deposit sites. To identify the type of microglia recruited in the amyloid plaques, we subclustered the entire microglial population and identified a subcluster particularly overrepresented in brain regions with amyloid plaques (Figure 4D right). Interestingly, this subcluster is also identified in WT sections, close to the external border of the brain (Figure 4D,E). This position suggests that they could be part of the border-associated macrophages (BAM) population. This subcluster is characterized by a higher expression of inflammation markers Igf1 and Igfbp5. On the other hand, the other microglia present in the cortex of APP mice express markers promoting cell survival and reduction of neuroinflammation (Figure 4F). Interestingly, microglia are almost completely absent from the striatum and BAM are not recruited to the hypothalamic region in the APP sections. (Figure 4D).

**Figure 4.**
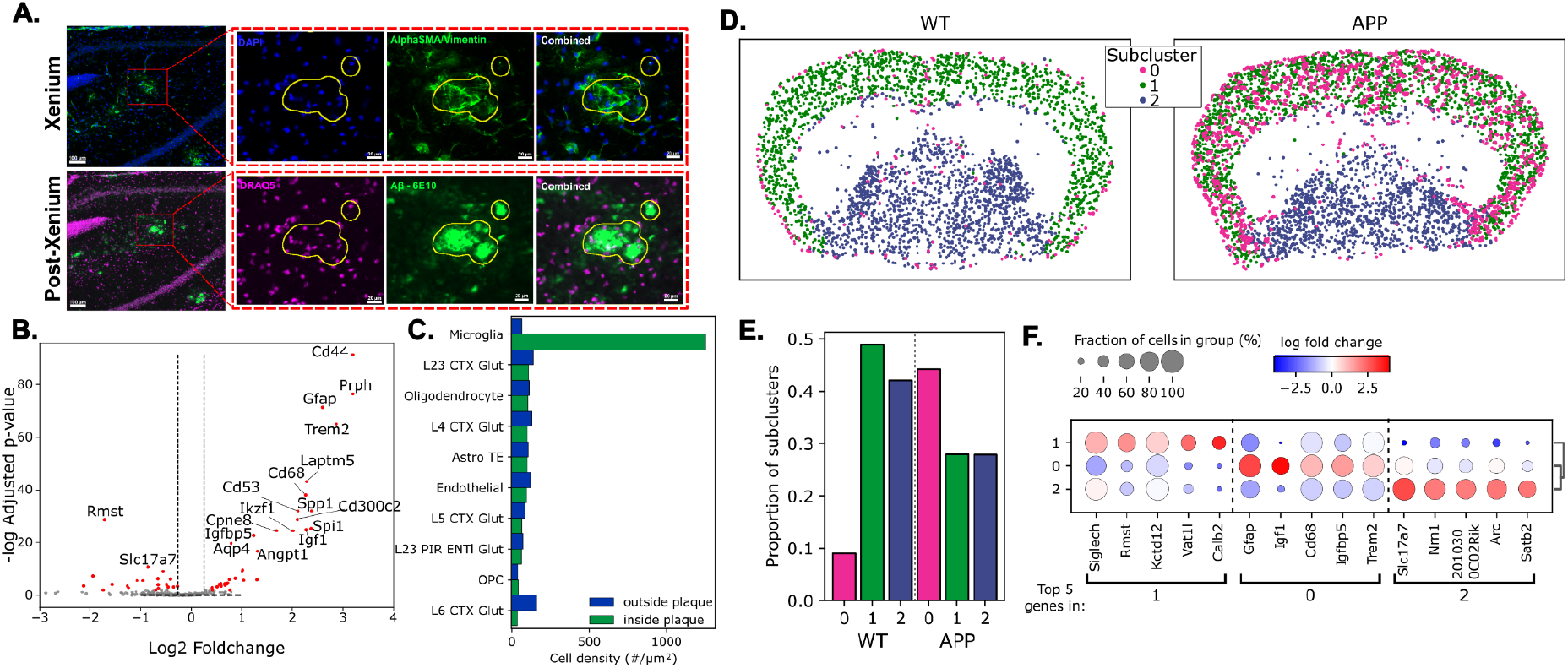
Recruitment of border associated macrophages within amyloid plaques. A. Example of amyloid plaques (yellow contour) identified by immunostaining (6E10, lower panel) correlating with alphaSMA staining (Xenium cell staining). B. Volcano plot of differentially expressed genes within amyloid plaques. The red dots correspond to genes significantly different. C. Density of cortical cells identified outside and inside the border of the amyloid plaques. D. Example of distribution of subclusters of microglia in WT and APP sections. The pink dots correspond to the border associated macrophages. E. Proportion of each subcluster in WT and APP sections. F. Dotplot representing the top 5 differently expressed genes in each subcluster of microglia. Dot size represents the percentage of cells expressing the gene in that subcluster and dot color represents the fold-change compared to the other two subclusters.

### 5. The suprachiasmatic nucleus shows important changes in gene expression without signs of inflammation

We then examined the SCN for similar markers of AD. First, we did not observe changes in the cell population of the SCN (Figure 5A). The main cell population of the SCN is GABAergic neurons, associated with a small number of astrocytes, oligodendrocytes and endothelial cells. Only a few microglia were accounted for across all samples. Non-neuronal cells did not show any DEG. However, SCN neurons presented a total of 26 DEG between WT and APP samples, including 14 upregulated genes (Figure 5B). Interestingly, most of the frequently DEG related to immune function found upregulated in the cortex are not significantly different in the SCN. Instead, the upregulated genes are associated with synapse organization and plasticity, development, growth, and cell-cell adhesion. The downregulated genes are related to synapse function (response to calcium and vesicle trafficking), and to cell morphology.

**Figure 5.**
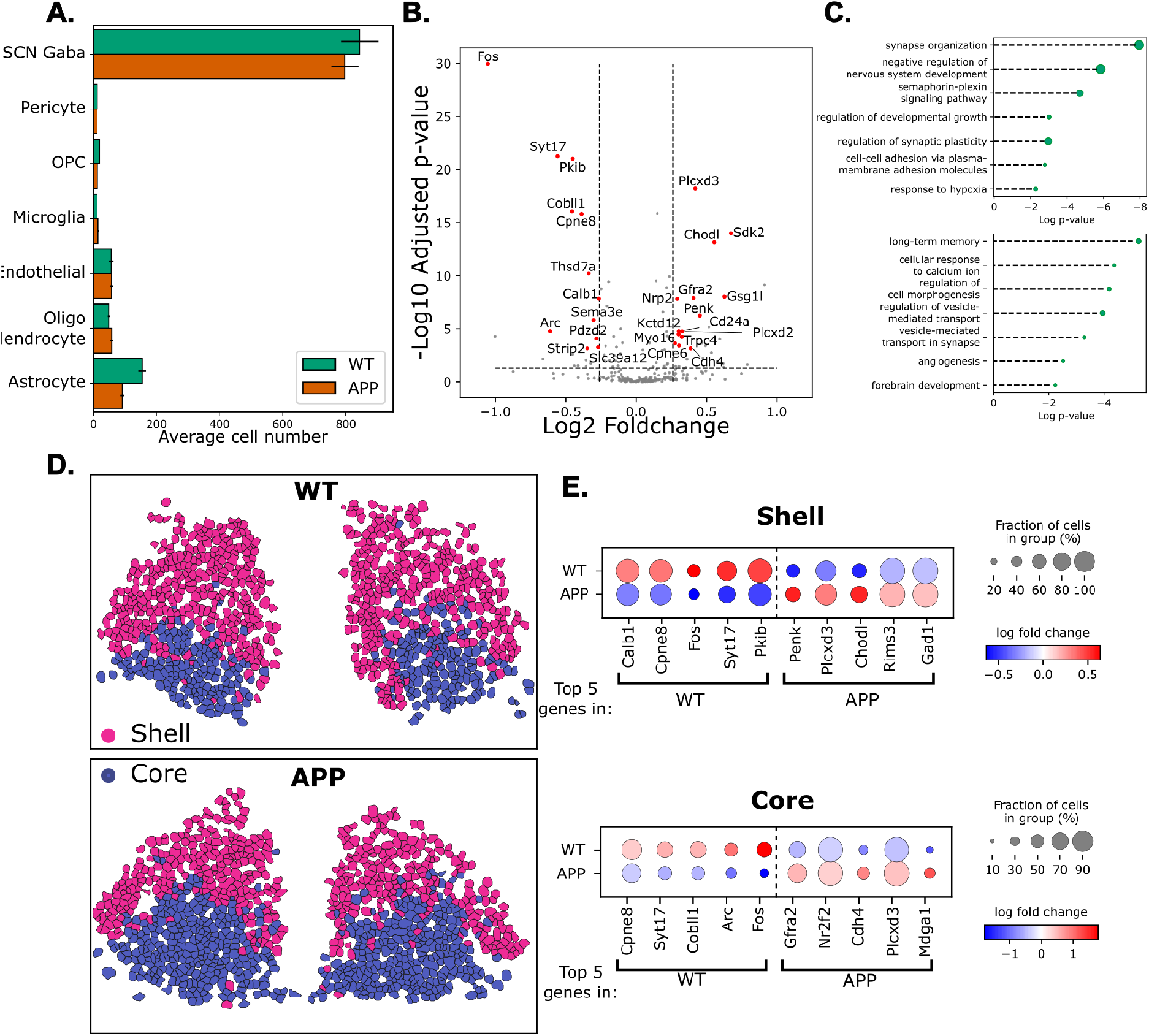
SCN neurons in WT and APP mice. (A) Average cell numbers for major brain cell types in WT and APP SCN. (B) Volcano plot of differentially expressed genes in SCN GABA neurons (APP vs. WT). The red dots correspond to genes significantly different. (C) Gene Ontology enrichment of upregulated (top) and downregulated (bottom) genes in APP SCN neurons. (D) Subclustering of SCN neurons allows identification of the shell (pink) and core (blue) regions of the SCN. (E) Dotplot representing the top 5 differently expressed genes in each subcluster of SCN neurons. Dot size represents the percentage of cells expressing the gene in that subcluster and dot color represents the fold-change compared to the other subcluster.

The SCN is known to be composed of two main regions, the ventral core and the dorsal shell, mainly composed of neurons expressing the neuropeptides VIP and AVP, respectively. Our previous studies showed region-specific organization and connectome (Calligaro et al., 2023). To see if the changes induced by AD are similarly region-specific, we subclustered the SCN Gabaergic neurons and were able to distinguish the two regions, in both WT and APP samples (Figure 5D). By examining the changes in gene expression in both regions, we observed that, despite both core and shell having differences between WT and APP, the top DEG had larger fold changes in the core region compared to the shell region of the SCN (Figure 5E). In particular, the reduction of the expression of the marker of activity *Fos* in the core region of APP samples could be due to a reduction of synaptic input. The core region of the SCN contains VIP neurons that are known to receive direct synaptic input from the retina inducing the expression of *Fos*. These results suggest that, despite not showing the classical signs of neuroinflammation observed around amyloid plaques, or the recruitment of immune cells, the SCN function is affected in this AD model. Thus, we next went to investigate the consequences of AD on the ultrastructures and connectome of the core SCN.

### 6. The cell content of SCN neurons changes with AD progression

To understand the cellular origin of the observed changes in circadian-related functions and in non-visual photoreception, we investigated whether the progression of AD affects the SCN connectome. To do so, serial-blockface scanning electron microscopy (SBEM) allows accessing the SCN network at the highest resolution and was previously used to establish the detailed input of mRGCs into the SCN network in WT mice (Calligaro et al., 2023; Kim et al., 2019) and in the study of the consequences of neurodegenerative diseases (Panes et al., 2023). We obtained samples of 3- and 8-month-old APP/PS1 mice (respectively, 3M and 8M) and collected image volumes of the ventral SCN (core SCN) and the olivary pretectal nucleus (OPN), another brain region targeted by mRGCs. We then manually reconstructed the connectome and the intracellular content of the neurons.

We first counted the number of neurons, glial cells, and endothelial cells present in each volume. We observed a reduced density of neurons and an increased density of glia and endothelial cells (Figure 6A), which is consistent with the neuronal loss and gliosis expected in AD. We then quantified the contents of the soma to determine if AD is associated with somatic changes in the SCN. We reconstructed the nuclei of each soma but observed no significant change in their volume between the 3M and 8M volumes (3M =316.79 ± 8.70µm3, 8M = 365.01 ± 21.02 µm3, p = 0.05, Figure 6B). However, we observed a change in their shape. In the 3M volume, all neuronal nuclei presented a complex shape with multiple infolding patterns (Figure 6C). In the 8M volume, we observed a mix of infolded and completely spherical nuclei (Figure 6C).

**Figure 6.**
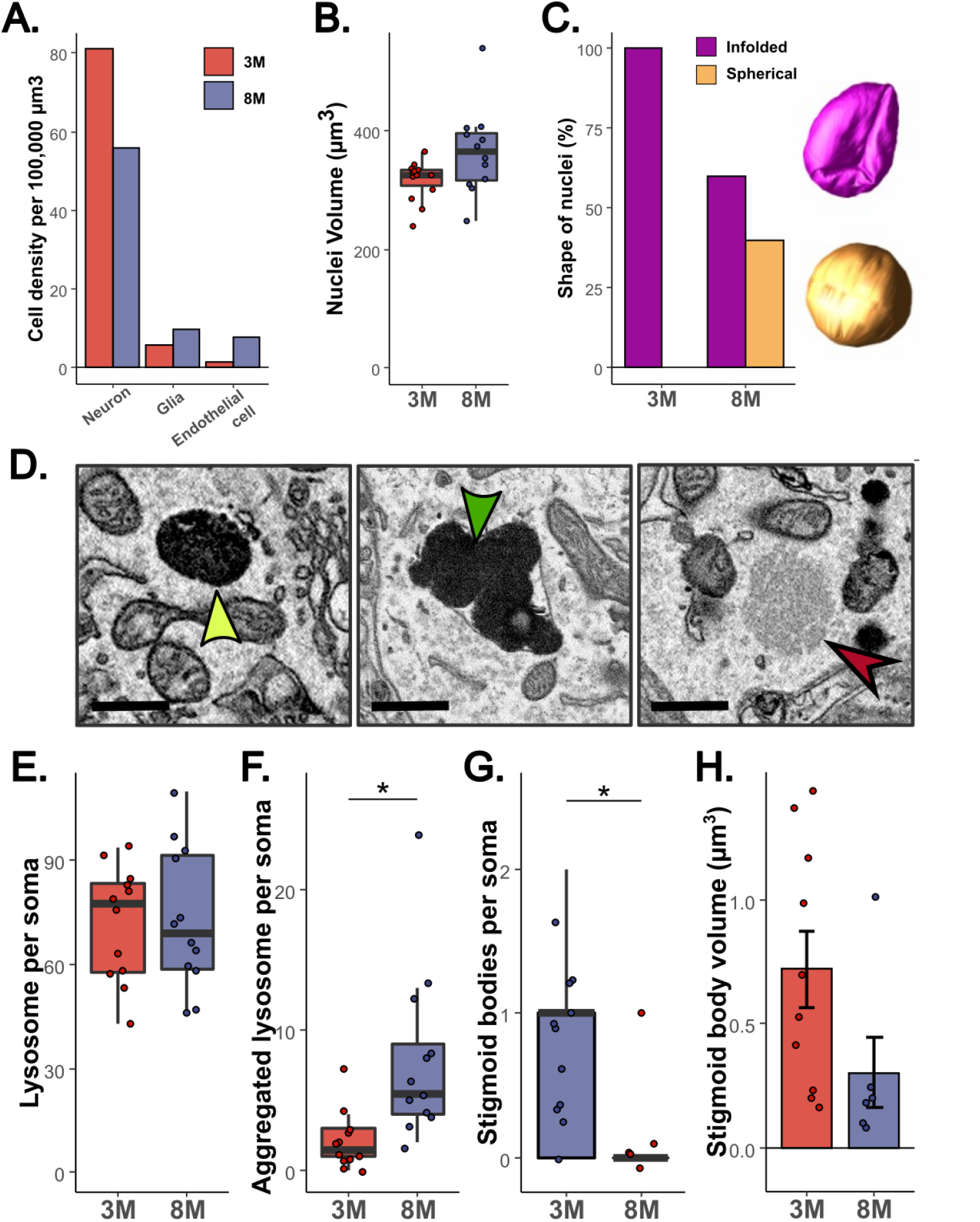
Cell somas undergo significant alterations in their cellular contents. (A) Cell densities in the SCN image volumes. (B) Volumes of nuclei in identified neurons. (C) Proportion of neuronal nucleus separated by shape. (D) Representative examples of a lysosome (left), an aggregated lysosome (middle) and a stigmoid body (right). Lysosomes were subcategorized into unaggregated and aggregated lysosomes depending on whether the lysosomes were clustered together. Scale bar = 500 nm. Quantification of (E) lysosomes, aggregated lysosomes (F), stigmoid bodies (G), and measure of stigmoid bodies volume (H). Results are expressed as mean +/- SEM. Statistical analysis done using Wilcoxon Signed Rank test, * = p < 0.05.

While we found a similar number of single lysosomes (Figure 6D, left) in both SCN (3M = 71.92 ± 4.76 lysosomes/soma, 8M = 73.08 ± 5.92 lysosomes/soma, p = 0.88, Figure 1E), the 8M had a significantly higher number of aggregated lysosomes (Figure 6D, middle) in its somas, which can be associated with neurodegenerative diseases (3M = 2.08 ± 0.16 aggregated lysosomes/soma, 8M = 7.83 ± 0.51 aggregated lysosomes/soma, p < 0.001, Figure 6F). We observed a decrease in the number of stigmoid bodies (Figure 3D, right), a cytoplasmic structure resembling a nucleolus, in the somas of the 8M SCN (3M = 0.67 ± 0.05 stigmoid bodies/soma, 8M = 0.08 ± 0.02 stigmoid bodies/soma, p < 0.05, Figure 6G). These stigmoid bodies tended to be smaller in the 8M SCN (3M = 0.72 ± 0.16 µm3, 8M = 0.3 ± 0.14 µm3, p = 0.055, Figure 6H).

### 7. AD progression affects pre- and post-synaptic structures

Out of 100 randomly selected and fully reconstructed axons (Supplementary Videos 2,3), we observed an increase in bouton frequency in individual axons in the 8M SCN (3M = 7.09 +/- 0.42 boutons/100 µm; 8M = 9.81 +/- 0.58 boutons/100 µm; p-value < 0.001; Figure 7B), without an increase in the total bouton density (3M = 13.95 +/- 1.52 µm3; 8M = 13.49 +/- 2.15 µm3; p-value = 0.88; Figure 7C). This apparent paradox could be explained by a reduction of the total number of neurons, with the remaining ones forming more boutons as a compensation mechanism. Some axons can have dendritic intrusions invaginating inside their boutons (Figure 7A). These structures increase the exchange surface between the pre- and post-synaptic elements. We previously demonstrated that the large majority of axons originating from the retina do not have dendritic intrusions in their boutons (Calligaro et al., 2023). Based on this classification, we observed a slight increase in the number of boutons with dendritic intrusions in the 8-month SCN (Figure 7D). The boutons with intrusions are generally larger, have more synaptic partners, and more mitochondria compared to boutons without intrusions (Figure 7E-G). The boutons without intrusions presented a reduction in the size of the boutons in the 8M, while synaptic density and mitochondrial volume remained stable between 3 and 8 months.

**Figure 7.**
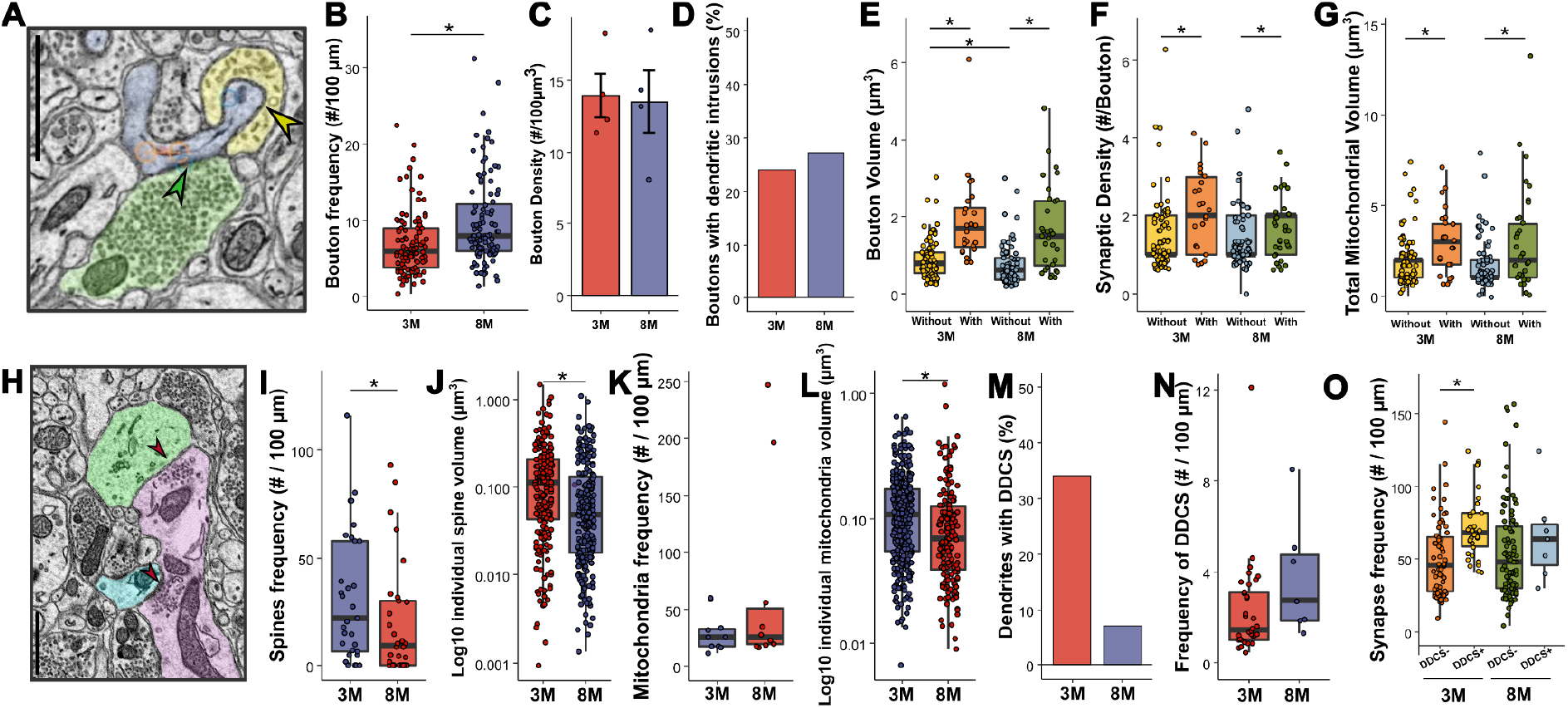
AD progression affects pre- and post-synaptic structures. (A) Representative examples of synaptic contacts between a dendrite (blue) and two axons (green and yellow), forming two types of boutons found in the SCN: boutons without (green arrowhead) and with dendritic intrusions (yellow arrowhead). Scale bar = 1000 nm. Quantification of linear bouton frequency (B), density (C), and proportion of boutons containing dendritic intrusions (D). Quantification of bouton volume (E), synaptic density (F), and mitochondrial volume (G) after segmentation. (H) Representative illustration of dendro-dendritic chemical synapses (DDCS, red arrowheads) between SCN dendrites. Scale bar = 1000 nm. Quantification of dendritic spines (I, J) and mitochondria (K, L) frequency and volume. (M) Percentage of dendrites forming DDCS. (N) Frequency of DDCS in DDCS-positive dendrites. (O) Synapse frequency in DDCS-negative and DDCS-positive dendrites. Statistical analysis done using the Wilcoxon Signed Rank test, * = p < 0.05.

To examine the post-synaptic elements of the connectome, we fully segmented 100 random dendrites, their contents, and identified all synapses. The dendrites of the 8M SCN present a reduced spine linear frequency (3M = 31.9 +/- 5.4 spines/100 µm; 8M = 19.9 +/- 4.7 spines/100 µm; p-value <0.05, Figure 7I) combined to a reduced volume of individual spines (3M = 0.16 +/- 0.01 µm3; 8M = 0.10 +/- 0.01 µm3; p-value <0.001, Figure 7J). The mitochondria in 8M dendrites were also smaller (3M = 0.1317 +/- 0.0056 µm3; 8M = 0.1126 +/- 0.0108 µm3; p-value < 0.001, Figure 7L) despite no change in their frequency (3M = 29.5 +/- 5.3 mitochondria/100 µm; 8M = 65.6 +/- 26.5 mitochondria/100 µm; p-value = 0.63, Figure 7K). The SCN contains a network of dendrites that form dendrodendritic chemical synapses (DDCS) and receive denser synaptic input than dendrites without DDCS (Calligaro et al., 2023; Kim et al., 2019). A smaller proportion of dendrites with DDCS was found in 8M SCN compared to 3M SCN (3M = 37%; 8M = 7%, Figure 7M). Their frequencies were similar, however (3M = 2.29 +/- 0.36 DDCS/µm; 8M = 3.7 +/- 0.96 DDCS/µm; p-value = 0.1143; Figure 7N). As documented previously, we observed a higher synaptic input onto dendrites with DDCS (DDCS+) compared to dendrites without DDCS (DDCS-) in 3M SCN (DDCS- = 50.90 +/- 3,28 synapse/100 µm; DDCS+ = 72.16 +/- 3.77 synapses/100 µm; p-value <0.001, Figure 7O), but this difference in synaptic input disappears in 8M SCN (DDCS- = 54.75 +/- 3,38 synapse/100 µm; DDCS+ = 65.45 +/- 11.59 synapses/100 µm; p-value = 0.17, Figure 7O). This could indicate a reduction of DDCS network innervation and potentially of retina-originated innervation.

Taken together, these results could suggest a limited impact of AD on the SCN connectome, with an overall relatively stable density of synapses. There are, however, signs of a reduction in retinal axon inputs. Additionally, the results could be impacted by a selection bias, as the axons and dendrites selected for reconstruction were chosen randomly using a grid pattern and morphological criteria (see Methods). We thus selected “normal-looking” axons and dendrites. However, after a systematic segmentation of the dendrites of neurons whose soma were within the volume, we identified strong dystrophy signs in 25% of 8M SCN dendrites (Figure 8A-D), including unusual shapes, intertwining of adjacent dendrites, invagination of axons within the soma, and an absence of any synapse on all these segments. Similarly, we identified multiple swollen axons, filled with debris or clusters of damaged mitochondria within what appears to be a membrane, possibly the result of unsuccessful recycling (Figure 8E-G). We also identified microglia filled with aggregated lysosomes and lipid droplets (Figure 8H). No sign of neurodegeneration was observed in the olivary pretectal nucleus (OPN) samples from the same mice (Supplementary Figure 3).

**Figure 8.**
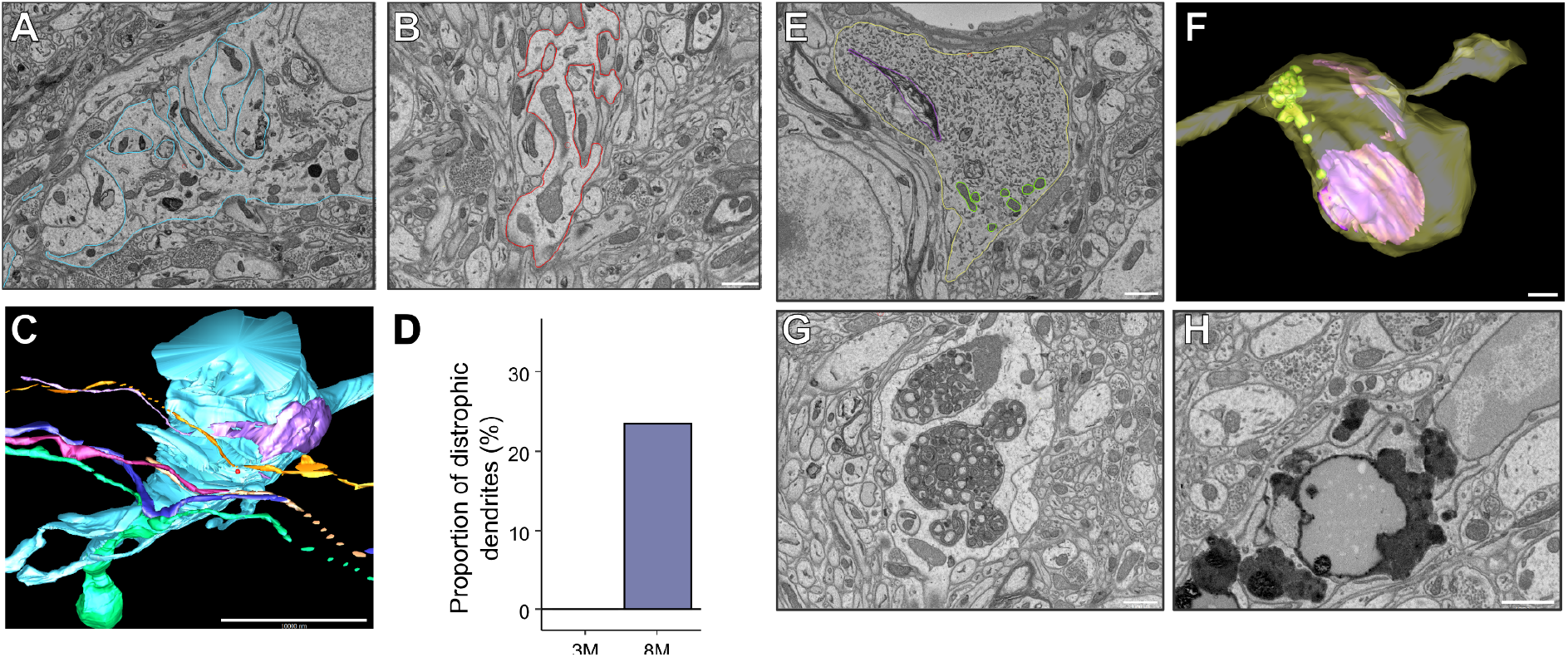
The SCN of an 8M mouse presents multiple signs of neurodegeneration. A,B. Representative images of dystrophic dendrites and soma. C. 3D reconstruction of the dendrite from panel A with surrounding axons. D. Proportion of dendrites presenting dystrophic dendrites. E,F. EM image (E) and 3D reconstruction (F) of a swollen axonic bouton filled with debris (purple) and mitochondria (yellow). G. Cluster of dystrophic mitochondria surrounded by a membrane within an axon. H. Accumulation of aggregated lysosomes and lipid droplets inside the soma of a microglia.

## Discussion

We presented here a multi-scale, in-depth investigation of the consequences of early Alzheimer’s disease on brain gene expression, the SCN connectome, light photoreception, and downstream SCN-controlled functions. In this work, we characterized early-onset circadian symptoms, including alterations in sleep architecture, locomotor activity, core body temperature, and phase shifting. Even at an early stage of the disease, we identified multiple signs of progression, indicating that circadian dysfunctions emerge early and evolve over time.

The effect of AD on sleep regulation in mice models is strain specific (Kent et al., 2018), but consistently shows alteration of their sleep patterns (Duncan et al., 2026; Kent et al., 2018; Van Erum et al., 2018; Whittaker et al., 2023). The classification of subclusters of NREM sleep allows to access in more detail the differences happening close to the transitions from dark to light, and suggests a reduced sleep efficiency, as observed in patients with AD (Ju et al., 2013). Studies in rodents report a close, bilateral link between the functioning of the SCN and the regulation of sleep patterns, in particular REM sleep (Wurts and Edgar, 2000; Deboer et al., 2003; Van Erum et al., 2018). In patients, the number of VIP-expressing neurons of SCN decreases with disease progression, possibly impacting its function in regulating circadian rhythms (Swaab et al., 1985; Zhou et al., 1995).

Consistent with this hypothesis, we observed substantial alterations in the SCN connectome in our mouse model of AD. The most prominent structural change is a marked decrease in dendrites with DDCS. This dendritic network has a central role in synaptic integration, as it receives the majority of synaptic inputs in the healthy mouse SCN (Calligaro et al., 2023; Kim et al., 2019). Thus, its disruption likely impairs the integration and processing of photic and non-photic signals. Moreover, these effects seem to be limited to the SCN at this age and stage of the disease as the second retino-recipient region we examined, the OPN, do not present signs of dystrophy. To note, the OPN receives fewer mRGC axons compared to the SCN (Hattar et al., 2002), and receives mainly non-M1 mRGC subtypes, while the SCN receives exclusively M1 subtypes (Aranda and Schmidt, 2020), and presents a different organization of synaptic input (Kim et al., 2019). The OPN could be affected by AD at later stages of the disease, which could lead to the pupillary light reflex deficits observed in other AD mouse models (Recio et al., 2026) and patients (Oh et al., 2019).

At the molecular level, gene expression analysis of the SCN in APP/PS1 mice reveals significant changes despite the absence of detectable amyloid-beta accumulation at any age. Notably, the SCN does not exhibit AD-associated markers of inflammation but instead shows strong alterations in genes related to cell morphology and synapse organization. Together, these findings indicate that the central clock undergoes substantial structural and functional remodeling early in the disease. This remodeling provides a mechanistic basis for the circadian-related symptoms observed in Alzheimer’s disease.

The specific origin of SCN changes in older APP/PS1 mice is unclear but a thread of evidence leads to a relation between light photoreception and AD progression. Melanopsin photoreception is strongly and specifically affected in AD patients (La Morgia et al., 2016, 2017; Oh et al., 2019; La Morgia et al., 2023; Sanda et al., 2026) and in mouse models (Recio et al., 2026). Additionally, dim light at night, a form of light pollution, can increase the plaque burden in the cortex and hippocampus of AD mouse models (Duncan et al., 2026). We thus sought to investigate the evolution of mRGC functions with AD progression. Our finding that in young APP/PS1 mice, the mRGCs presented a hyperactive response to light while older APP/PS1 mice presented a reduced response drove us to the hypothesis that this early hyperactivity induces excitotoxic effects in downstream targets, particularly within the SCN. It could explain the observed deficits in absence of the usual markers of AD pathology, microglia recruitment, inflammation and amyloid deposits.

Over the past decades, research has primarily focused on amyloid-beta oligomers and deposits, their local effects, and strategies for their clearance. As a result, brain regions with prominent plaque accumulation, such as the cortex, hippocampus, and amygdala, have been extensively studied, and the other regions such as the hypothalamus were less investigated while potentially holding crucial information to understand the intertwined interactions between brain regions affected by AD. Spatial single-cell transcriptomics have been a powerful tool to investigate these changes in gene expression in brain tissue (Zhang et al., 2024), both at a local scale (e.g. around amyloid plaques) as well as at a more global scale. Nonetheless, studies have been focusing on specific cell-types (astrocytes and microglia (Sadick et al., 2022; van Olst et al., 2025; Wei et al., 2025; Liu et al., 2026)). Here, we generated a larger view of the consequences of AD on gene expression across multiple brain regions, allowing us to identify the region-specific effects of AD.

In addition, we introduced a method to exploit the full potential of imaging-based single-cell spatial transcriptomics. By analyzing transcripts not assigned to segmented cell soma, we identified extrasomatic RNA populations and confirmed previous observations of local translation within neurites, particularly in proximity to synapses. This approach provides additional spatial information that is not captured by standard cell-soma-based analyses. Using this method, we detected an increased number of inflammatory markers in the cortex beyond amyloid plaques, whereas the hypothalamus, including the SCN, exhibits a distinct molecular profile. This contrast highlights once more region-specific disease mechanisms, with inflammation dominating in plaque-rich cortical regions and structural and synaptic alterations prevailing in the hypothalamus, supporting the results obtained using somatic transcripts. To note, during the preparation of this manuscript, a preprint describing a similar approach to investigate extrasomatic transcripts, referred to as unassigned RNA (uRNA), was published on bioRxiv (Salas et al., 2025).

## Supplementary figures

**Supplementary figure 1:**
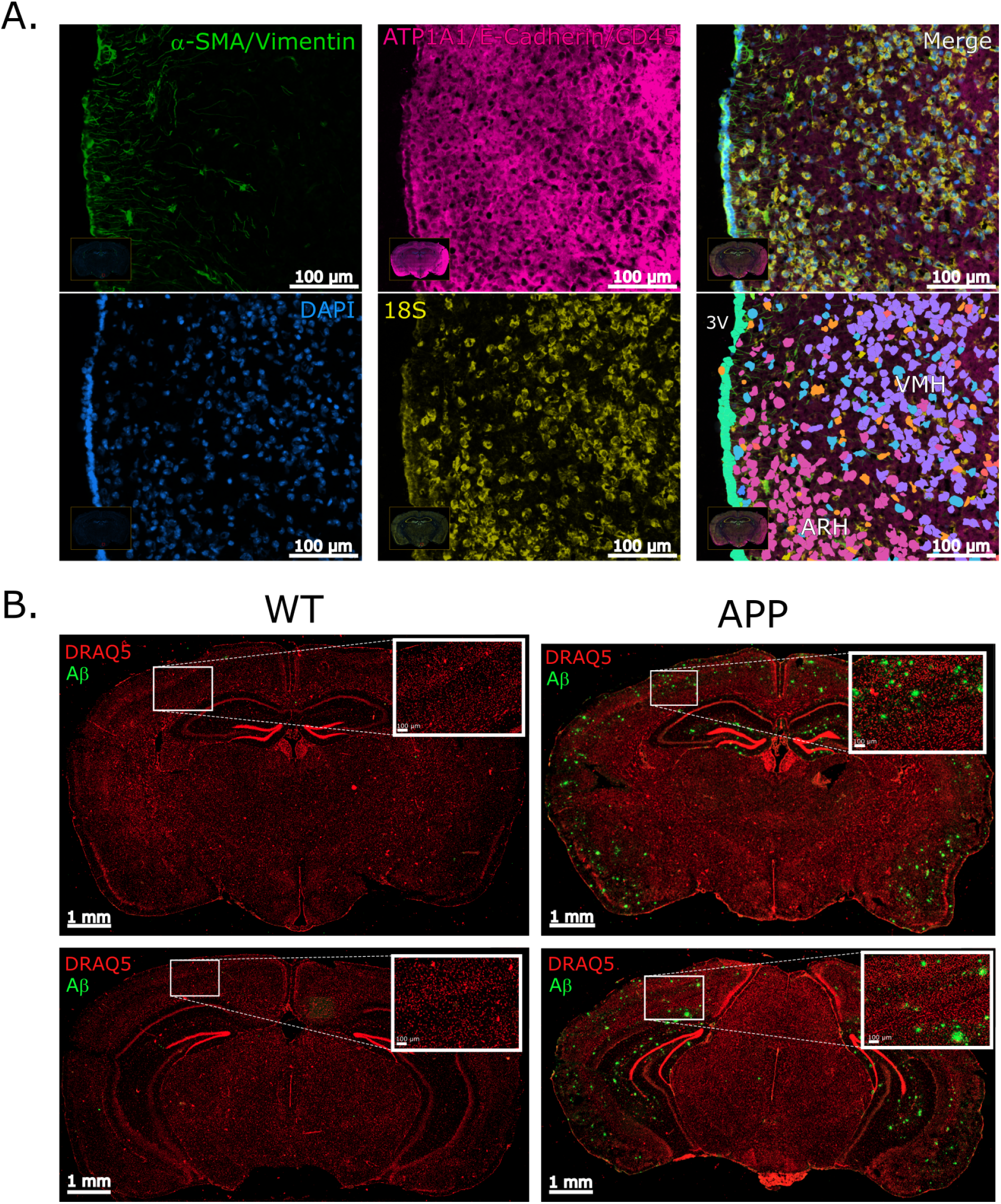
A. Cell markers used for cell segmentation by Xenium Explorer. Lower right panel shows cell segmentation with each color corresponding to a cell type. DAPI = 4′,6-diamidino-2-phenylindole, α-SMA = alpha-smooth muscle actin; 3V = third ventricle, ARH = Arcuate nucleus, VMH = Ventromedial nucleus of the hypothalamus. Scale bar = 100 µm. B. Examples of Amyloid-beta stainings in WT (left) and APP/PS1 (right) mice. Red = DRAQ5 (nucleus); green = Amyloid-beta (scale bar = 1mm). Inserts are zoomed in cortex (scale bar = 100 µm)

**Supplementary Figure 2:**
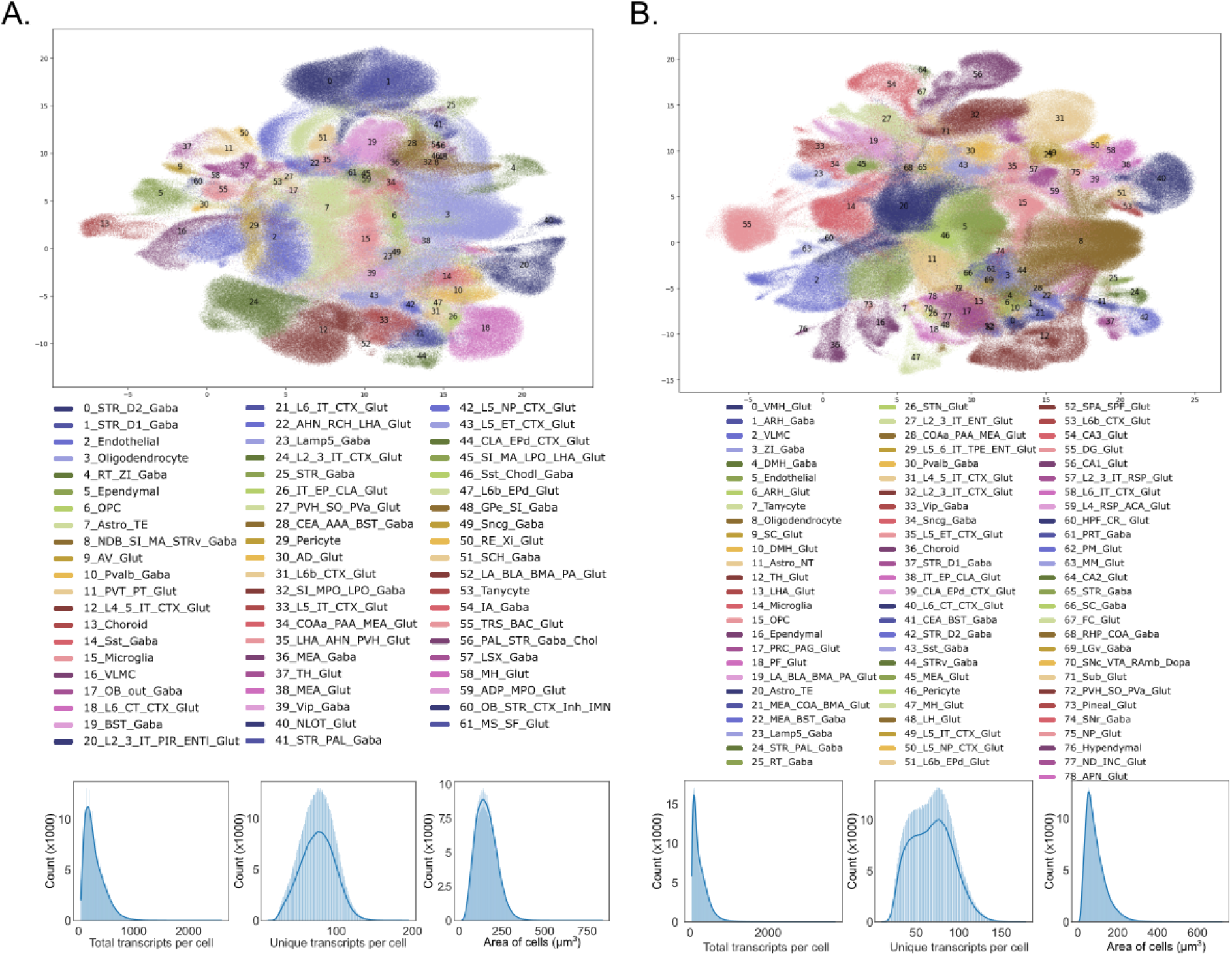
Annotated UMAP (top) and cell metrics (bottom) of spatial transcriptomic samples from R0 (A) and R1 (B) experiments.

**Supplementary Figure 3:**
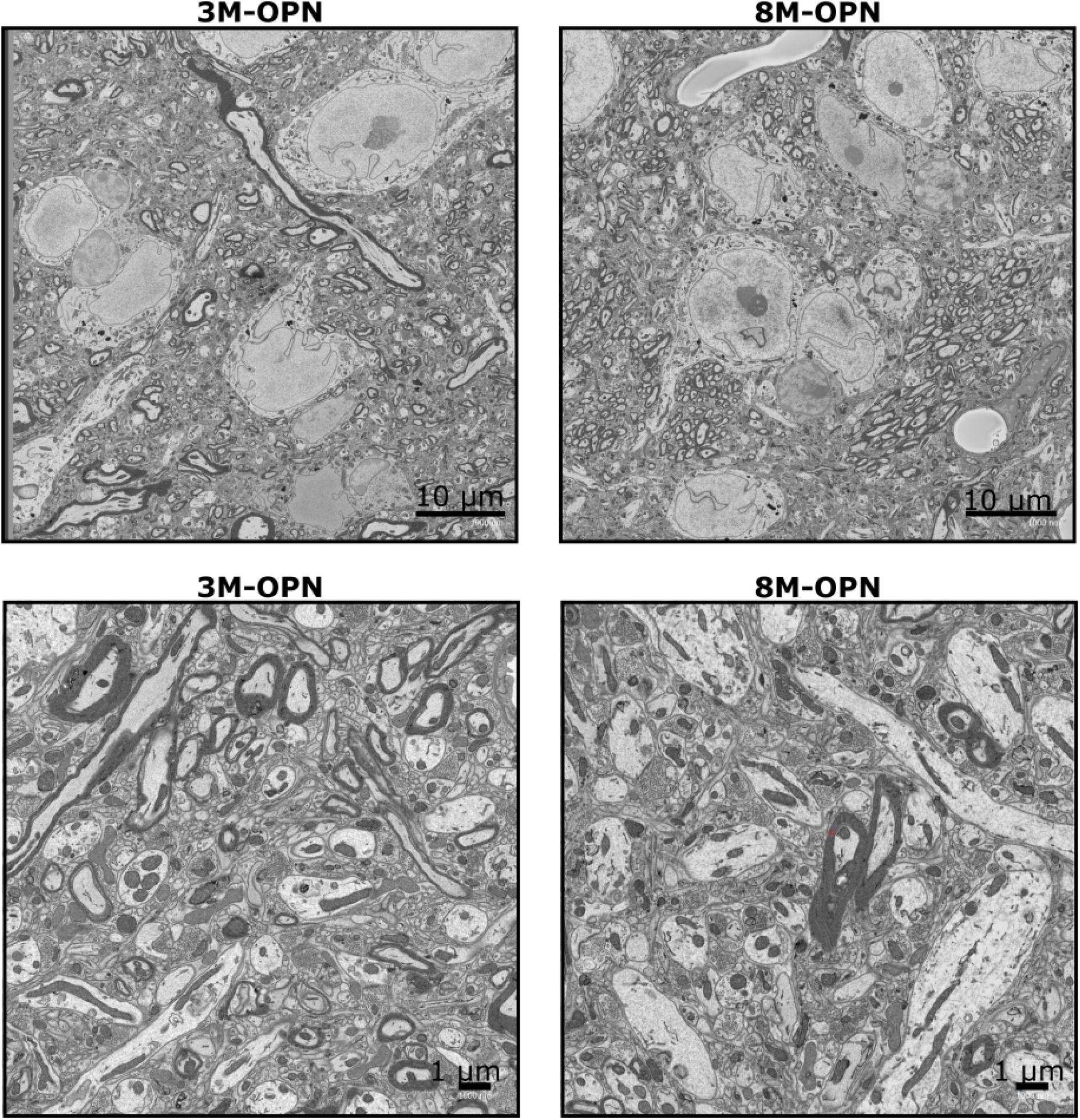
Representative images of zoomed out (top) and zoomed in (bottom) image volume of 3M and 8M OPN.

---

[Video 1: examples of structures in SBEM](https://drive.google.com/open?id=1cdGIPPGO7vzI5y7NdQgkZmKJ4SPOtQFC) Examples of glial processes, cell soma, dendrites, and axons, in order.

[Video 2: Visualization of axons, dendrites and synapses in 3M SCN](https://drive.google.com/file/d/1-9AFO9T7nMnvufydUWsbEEgcM1GMu-Sd/view?usp=drive_link)

[Video 3: Visualization of axons, dendrites and synapses in 8M SCN](https://drive.google.com/file/d/1ttSgRrP2M5J-NJ80595-Ym9UaP2p9i9q/view?usp=drive_link)

## Acknowledgments

We thank Matthew Le, Caitlyn Kim, and Marianne Aquino for their help with SBEM segmentation, and Vince Rothenberg for his help on writing custom scripts for SBEM and spatial transcriptomic analyses. Rita and Richard Atkinson Endowed Chair fund to SP

## Contributions

HC, SP, ME, MTYL designed the experiments. HC, KYK, KN, IL, LvR collected the data. HC, BK, KN, MTYL, RR, LvR, AG, SP, ME analyzed the data. HC prepared the first draft of the manuscript. All authors reviewed and edited the final manuscript.

